# Development of an Antiepileptogenesis Drug Screening Platform: Effects of Everolimus and Phenobarbital

**DOI:** 10.1101/2020.12.14.422712

**Authors:** Melissa Barker-Haliski, Kevin Knox, Dannielle Zierath, Zachery Koneval, Cameron Metcalf, Karen S. Wilcox, H. Steve White

## Abstract

**Objective:** The kainic acid (KA)-induced status epilepticus (SE) model in rats is an etiologically-relevant animal model of epileptogenesis. Just as in patients, who develop temporal lobe epilepsy (TLE) following SE, this rat model of KA-induced SE very closely recapitulates many of the clinical and pathological characteristics of human TLE that arise following SE or another neurological insult. Spontaneous recurrent seizures (SRS) in TLE can present after a latent period following a neurological insult (TBI, SE event, viral infection, etc.). Moreover, this rat model of TLE is ideally suited for preclinical studies to evaluate the long-term process of epileptogenesis and screen putative disease-modifying/antiepileptogenic agents. This report details the pharmacological characterization and methodological refinement of a moderate-throughput drug screening program using the post KA-induced SE model of epileptogenesis in male Sprague Dawley rats to identify potential agents that may prevent or modify the onset or severity of SRS. Specifically, we sought to demonstrate whether our protocol could prevent the development of SRS, or lead to a reduced frequency/severity of SRS.

**Methods:** Rats were administered everolimus (2-3 mg/kg, P.O. commencing at 1, 2, or 24-hrs after SE onset) or phenobarbital (60 mg/kg, beginning 1 hr after SE onset). The rats in all studies (n=12/treatment dose/study) were then followed intermittently by video-EEG monitoring; i.e., 2-weeks on/2-weeks off, 2-weeks on epochs to determine latency to onset of SRS, and disease burden following SRS onset.

**Results:** While there were no adverse side effects observed in any of our studies, no treatment conferred a significant disease modifying effect, nor did any agent prevent the presentation of SRS by 6 weeks post-SE onset.

**Conclusions:** While neither phenobarbital nor everolimus administered at several time points post-SE onset prevented the development of SRS, we herein demonstrate a moderate-throughput screen for potential antiepileptogenic agents in an etiologically-relevant rodent model of TLE.

**Key Points:**

- Disease-modifying therapies are needed to prevent or attenuate the burden of epilepsy in at-risk individuals.
- We report a moderate-throughput screening protocol to identify disease-modifying agents in a rat post-kainic acid status epilepticus model.
- Everolimus was administered at multiple time points post-status epilepticus with no effect on spontaneous seizures up to 6 weeks later.
- Repeated administration of phenobarbital also did not prevent the development of spontaneous recurrent seizures up to 6 weeks post SE.
- While we did not identify any effect of either agent, our approach provides a moderate-throughput screen for antiepileptogenesis.

## Introduction

One in 26 Americans will develop epilepsy in their lifetime. Despite over 30 antiseizure drugs (ASDs) available for the treatment of symptomatic seizures, no therapy is yet approved to prevent this disease in at-risk individuals,^1; 2^ substantiating the need for disease-modifying and/or antiepileptogenic agents. With this in mind, the National Institute of Neurological Disorders and Stroke (NINDS) refocused its long-standing Epilepsy Therapy Screening Program’s (ETSP) screening workflow to not only evaluate compounds that may treat the symptomatic seizures of drug-resistant epilepsy, but also identify agents that may potentially modify or altogether prevent the development of epilepsy as recommended by recent NINDS Advisory Council Working Group reviews of the ETSP.^3; 4^ Under the NINDS ETSP’s prime contract (HHSN271201600048C), the University of Utah has sub-contracted the University of Washington to evaluate the potential of putative disease-modifying agents in a post-status epilepticus (SE) model of temporal lobe epilepsy (TLE) in rats. One of the strengths of this subcontracted approach is that investigational compounds are subjected to evaluation by a large scientific team of experienced epilepsy investigators at the NINDS, University of Utah, and University of Washington. Further, this strategy employs an in-life testing protocol that models treatment paradigms in patients at-risk for acquired epilepsy.

Spontaneous recurrent seizures (SRS) in clinical TLE that develop after a latent period following a neurological insult (traumatic brain injury (TBI), SE, stroke, tumor, viral infection, etc.), are often focal impaired awareness seizures that generalize to tonic-clonic seizures, and are resistant to some FDA-approved ASDs.^5^ These features are reproduced in the rat systemic kainic acid (KA) model of post-SE induced TLE, which is characterized by SRS that results following a latent period. Post-KA SE rats exhibit marked reactive gliosis,^6^ neuroinflammation,^7^ and behavioral deficits.^8^ Moreover, the systemic low-dose KA-induced SE paradigm in rats is ideally suited for drug discovery to evaluate the impact of investigational agents on the development and severity of SRS.^9^

This study aimed to establish and validate an in-life screen using the repeated administration of the ASD, phenobarbital (PB), or the immunomodulator, everolimus (EVL) to rats immediately following the induction of SE. The primary study objective was to rigorously evaluate the extent to which pharmacological intervention after SE induction would modify the severity, and/or frequency of SRS up to 6-weeks later. This study describes the approach used to assess the antiepileptogenic potential of two investigational agents and several interventional time points in an effort to refine a screening protocol for the NINDS ETSP. Of note, this study was conducted in a blinded manner, with the University of Utah sequentially submitting each compound to investigators at the University of Washington (UW) as an unidentified compound for independent evaluation at the UW facility. PB was selected because previous studies have suggested the potential to prevent the development of post-traumatic brain injury-induced epilepsy in at-risk patients.^10-12^ EVL was selected because it is approved as a disease-modifying treatment for tuberous sclerosis complex (TSC),^13; 14^ a condition characterized by chronic seizures and epilepsy.^15; 16^ EVL is also a rapamycin derivative (rapalog) that inhibits mammalian target of rapamycin (mTOR) activation.^17^ It has been hypothesized that mTOR inhibition may exert antiepileptogenic effects in epilepsy and/or disease modification in TSC.^18-20^ The results of this blinded study demonstrates that neither PB nor escalating doses of EVL administered at discrete time points significantly prevented the development of SRS in our model. Despite the lack of antiepileptogenic effect with either agent using multiple study protocols, we herein establish the proof-of-concept that the NINDS-supported protocol conducted in the post-KA SE rat model of acquired epilepsy can rigorously evaluate potential antiepileptogenic agents in a moderate-throughput and unbiased manner.

## Method (*See Supplemental Files*)

### Results

#### Refinement of Testing Protocol

The primary goal of this study was to establish a testing strategy that would be appropriate to identify potential disease-modifying or antiepileptogenic agents. Thus, several time points of drug administration after the onset of sustained SE were evaluated (Figure 1) to ascertain the ability of PB (repeated injections beginning at a single time point) or EVL (repeated injections beginning at multiple time points after SE) to modify the onset and/or severity of SRS up to 6-weeks later. Using a sequential testing strategy, we evaluated the ability of EVL administered beginning 1-24 hours after KA-induced SE and repeatedly administered for 5-7 days post-SE onset. PB administration began 1 hour after SE onset, whereas EVL administration commenced at three time points – 1-, 2-, and 24-hours post-SE onset (Figure 1). The “original” testing protocol planned for the 24/7 vEEG monitoring for 0-14 days and 28-42 days post-SE; the “expedited” testing protocol planned for 24/7 time0synchronized video-EEG (vEEG) monitoring only from 28-42 days post-SE (Figure 1E). This expedited protocol resulted in significant cost savings [or efficiency] as we were able to focus our resources on those animals that survived the initial SE; i.e., stereotaxic EEG implant surgery was not conducted prior to SE induction, but three weeks post SE induction (see Figure 2 for details of different protocols).

**Figure 1.**
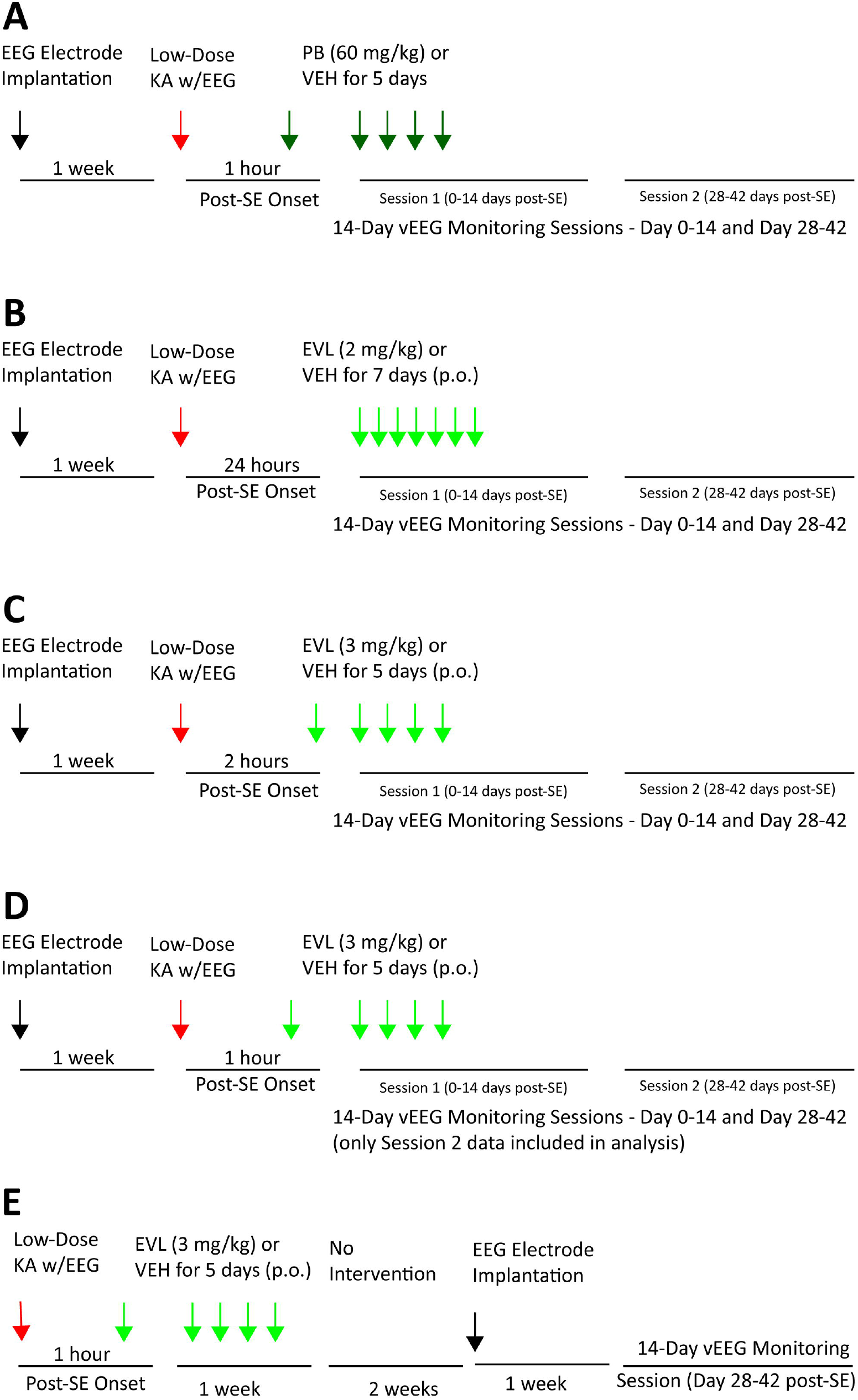
Original experimental protocol for the evaluation of the disease-modifying potential of A) phenobarbital (PB; 60 mg/kg, i.p.), B) everolimus (EVL; 2 mg/kg, p.o.), or C) EVL (3 mg/kg, p.o.) in male Sprague Dawley rats. Upon further refinement, the original protocol (D) was compared in a head-to-head study with an (E) expedited protocol to determine whether EVL (3 mg/kg, p.o.) would confer any disease-modifying effects against spontaneous recurrent seizure (SRS) severity or onset up to 6-weeks later. The original protocol design (A-D) included rats that were surgically implanted under ketamine/xylazine anesthesia 7-10 days prior to repeated low-dose administration of kainic acid (KA) to induce status epilepticus (SE) confirmed by video-EEG (vEEG) and sustained Racine stage 4/5 seizures for at least 30 minutes. Animals were randomly enrolled into either drug or respective vehicle (VEH) treatment groups and administered the first injection at 1 hour (A, D-E), 2 hour (B), or 24 hour (C) post SE induction. Rats were then intermittently monitored by continuous vEEG recording from 0-2 weeks and 4-6 weeks post-SE onset to determine latency to onset of SRS. In the expedited protocol (E), rats were implanted with EEG electrodes 7 days after KA SE and then only monitored from 4-6 weeks post-SE onset. In all studies, each investigational compound or VEH was administered by an experimenter blinded to treatment condition. Rats that survived the SE insult and investigational drug intervention period were monitored for the presence of spontaneous behavioral and electrographic seizures [or spontaneous behavioral seizures with electrographic correlate] up to 42 days post-SE insult. Video-EEG observed events were scored off-line by a trained investigator and confirmed by a secondary investigator, both of whom were blinded to experimental condition in all studies.

**Figure 2.**
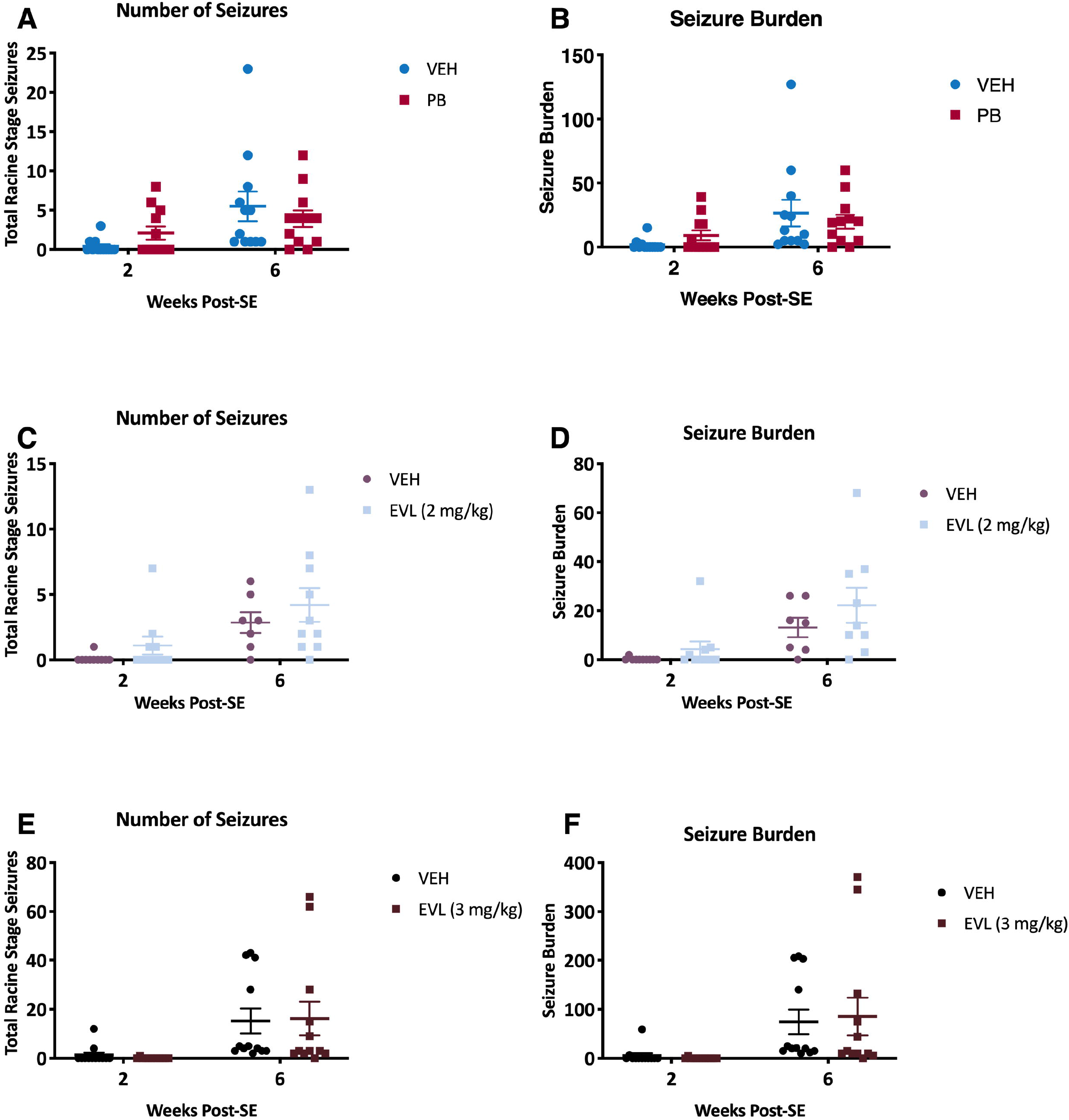
Recording session summary of the number of seizures and seizure burden during each 2-week recording epoch in the original protocol (two, 2-week recording sessions). There was no effect of phenobarbital (PB) on A) number of seizures during each recording session (0-2 weeks and 4-6 weeks post-SE) or B) average seizure burden. There was no effect of everolimus (EVL, 2 mg/kg) on C) the number of seizures during each recording session (0-2 weeks and 4-6 weeks post-SE) or D) average seizure burden. There was no effect of EVL (3 mg/kg) on E) the number of seizures during each recording session (0-2 weeks and 4-6 weeks post-SE) or F) average seizure burden.

#### Therapeutic intervention immediately after KA-induced SE did not significantly affect body weight change

KA-induced SE is typically associated with some body weight loss as a result of the unremitting seizure activity. There was no significant time x treatment interaction on body weight during the recovery period 4-7 days post-SE insult recovery period in any of the original testing protocol studies of this investigation. Specifically, there was no effect of repeated PB treatment on body weight change versus VEH-treated post-KA SE rats (Supplemental Figure 1A; F (5,86) = 143.0, p<0.0001). Additionally, EVL at either 2 mg/kg, p.o. (Supplemental Figure 1B; F (7, 140) = 0.2995, p<0.0001) or 3 mg/kg, p.o. (Supplemental Figure 1C; F (7, 126) = 3.320, p=0.0028) did not significantly affect body weight change versus their respective VEH-treated post-KA SE rats. In the blinded head-to-head comparison of the original versus expedited protocol study of EVL (3 mg/kg, p.o.), there was a significant effect of EVL treatment on body weight following SE (Supplemental Figure 2). Notably, there was no significant time x treatment interaction when EVL was administered after EEG implant (original protocol, Supplemental Figure 2A; F(4,104) = 1.587, p>0.18). However, when EVL was administered prior to EEG implant (expedited protocol, Supplemental Figure 2B; F(4,108)=4.241, p=0.0032) there was a significant effect of EVL treatment. Specifically, EVL lead to a reduction in body weight at Day 5 post-SE. Thus, treatment with PB and EVL for the 4-7 days following KA-induced SE was generally well tolerated and none of the investigational drug treatments lead to any notable worsening or improvements in body weight change versus their respective vehicle treatments when administered after the implantation of EEG probe (original protocol). However, administration of EVL prior to EEG implantation (expedited protocol) was associated with reduced body weight by day 5 post-SE.

#### Therapeutic intervention did not significantly reduce the number of seizures or seizure burden

Following the KA-induced SE insult, all rats in the study developed SRS within 6 weeks. Neither PB nor EVL treatment in any study significantly reduced the number of seizures that occurred during each of the 2-week monitoring sessions, and there was no effect of treatment on the seizure burden (i.e. modified Racine stage seizure severity x frequency of Racine stage) during these monitoring sessions. Specifically, there was a significant main effect of time on the total number of observed seizures for all studies, but no main effect of treatment or treatment x time interaction (Figure 2). Both PB and VEH-treated rats experienced a time-dependent increase in the total number of SRS (Figure 2A; F (1,22) = 7.968, p=0.0099) and seizure burden (i.e. seizure stage x frequency of events; Figure 2B; F (1, 22) = 7.323, p=0.0129). EVL (2 mg/kg) and VEH-treated rats also experienced a greater number of SRS (Figure 2C; F (1, 17) = 6.626, p=0.0197) and seizure burden (Figure 2D; F (1, 32) = 12.77, p=0.0011) with time post-SE. Finally, both EVL (3 mg/kg) and VEH-treated rats presented with a greater number of SRS (Figure 2E; F (1, 22) = 13.02, p=0.0016) and seizure burden (Figure 2F; F (1, 22) = 11.85, p=0.0023) with time post-SE. Similarly, no effect of EVL treatment started either at 1-, 2-, or 24-hours post-SE for 5 or 7 days duration was observed on either number of seizures nor seizure burden

Additionally, a head-to-head study of EVL (3 mg/kg) administered in the original or expedited protocols (Figure 3) demonstrated that there was no significant effect of protocol design on either the number of seizures (F (1, 39) = 0.08438, p=0.77) nor on seizure burden (F (1, 39) = 0.2168, p>0.64) in either treatment group. This suggests that the expedited protocol would be able to detect seizure events to a similar extent to that of the original protocol,

**Figure 3.**
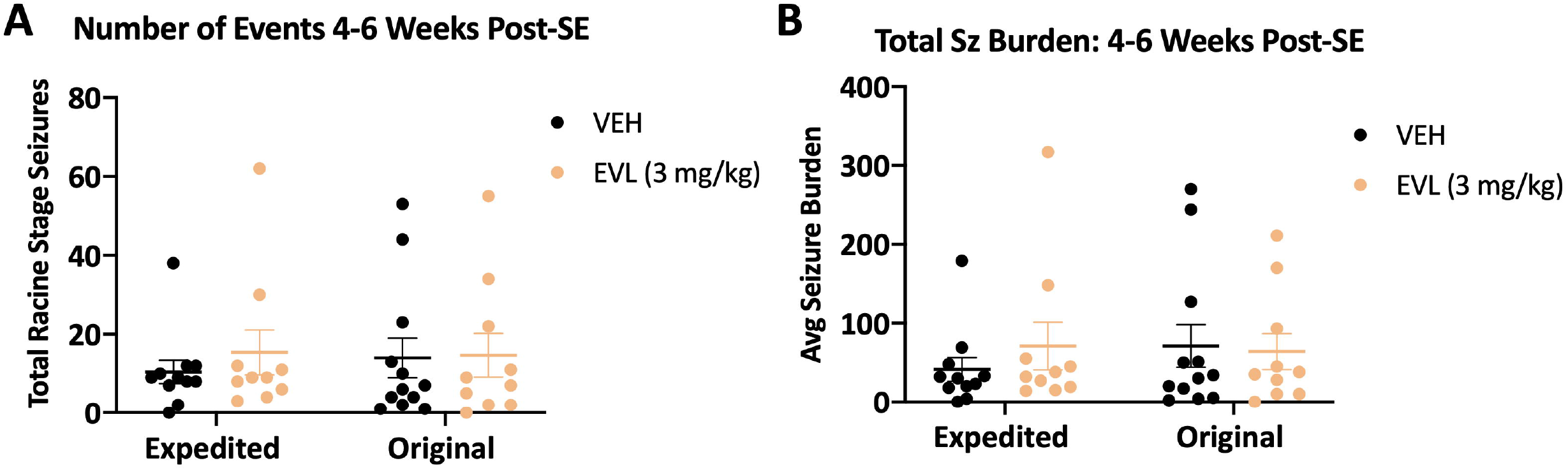
Recording session summary of the number of seizures and seizure burden during the 4-6 week recording epoch in the original (two, 2-week recording sessions) versus expedited protocol design. There was no effect of everolimus (EVL) on A) number of seizures during the 4-6 weeks post-SE recording session or B) average seizure burden.

#### The therapeutic interventions investigated did not significantly alter cumulative seizure burden up to 6 weeks post-SE

The cumulative seizure burden during the two, 2-week sessions within the 42-day recording session was designed to understand whether any intervention was able to alter the trajectory of disease progression over time. The study was not designed to determine the exact time of SRS onset. Cumulative seizure burden is designed to assess disease trajectory over time and includes the cumulative seizure severity x seizure frequency across all the observation days. PB administration did not significantly reduce the cumulative seizure burden versus VEH-treated control rats, although there was a general main effect of time post-SE (Figure 4A; F (55, 1210) = 15.36, p<0.0001). Similarly, EVL (2 mg/kg) treatment beginning 24 hours after SE onset did not alter the cumulative seizure burden versus VEH-treated control rats, although there was a general main effect of time post-SE (Figure 4C; F (51, 765) = 14.25, p<0.0001). Lastly, a higher dose of EVL (3 mg/kg) administered earlier after SE onset (2 hr; Figure 1C) did not significantly alter the cumulative seizure burden versus VEH-treated control rats, although there was a general increase in cumulative seizure burden with time post-SE (Figure 4E; F (49, 1029) = 9.221, p<0.0001). In the head-to-head study of the expedited protocol, administration of EVL did not significantly alter cumulative seizure burden in either study design (Figure 4G and 4I). There was, however, a general time-related increase in burden in both the original (F (1.141, 22.81) = 11.83, p=0.0016) and expedited study designs (F (1.275, 22.96) = 9.364, p=0.003). Thus, cumulative seizure burden was not significantly affected by any treatment versus the VEH-treated KA-SE rats within the same study group using this particular experimental paradigm.

**Figure 4.**
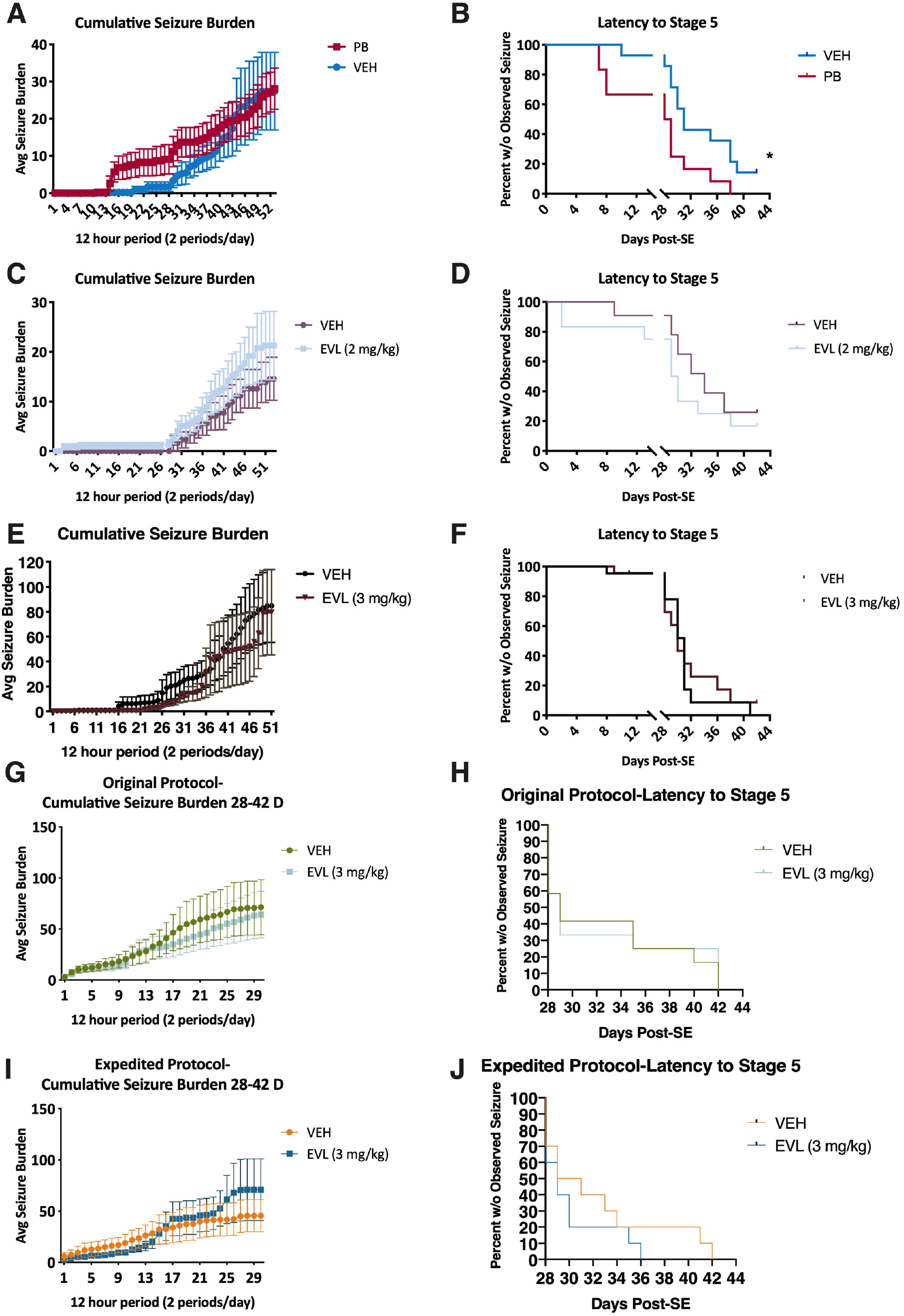
Cumulative seizure burden is the summation of all observed events during each of the two, 2-week-long recording sessions from 0-2 and 4-6 weeks post-SE (original protocol A, C, E). There was no significant effect of any investigational treatment A) phenobarbital, C) everolimus (EVL; 2 mg/kg), or E) EVL (3 mg/kg). B, D, F) The latency to stage 5 seizure was not significantly improved by any treatment condition for all rats up to 42 days post-SE in the original protocol. A) Administration of PB led to a significant reduction in the time until first observed Racine stage 5 seizure during the two, 2-week long monitoring sessions (X^2^=5.691, p=0.0171). While it is possible that the intermittent monitoring may have missed seizures that occurred during the 2-4 weeks post-SE using our original protocol, this analysis was conducted to further characterize disease progression in this novel drug screening model. In the head-to-head evaluation of EVL efficacy in the original versus expedited protocol (G-J), there was also no effect of EVL administration on cumulative seizure burden (G, I) nor on latency to stage 5 seizure (H, J). This head-to-head study demonstrates that the expedited protocol is valid for moderate-throughput disease modification studies and can demonstrate disease-modifying potential (or lack thereof) to an equivalent degree as the original, longer duration monitoring study of the original protocol.

#### The latency to stage 5 seizures was significantly reduced by PB treatment

The latency to the first observed spontaneous occurring Racine stage 5 seizure during the vEEG recording sessions was assessed to determine whether any intervention could effectively delay the onset of severe generalized seizures, an alternative metric of disease burden or disease course. Specifically, a compound that delays the time to onset of stage 5 seizures could suggest a disease-modifying effect. While it is possible that the intermittent sampling occurring 0-2- and 4-6-weeks post-SE onset (as used in the original study design) may have missed some seizures in the 2-4 weeks post-SE period, we did include this evaluation metric in an attempt to better characterize disease progression in our drug screening paradigm. Additionally, the expedited study design was similarly limited to only a 4-6 weeks post-SE session, but used to screen for potential agents that may effectively modify SRS severity or frequency. We herein demonstrate that PB, when administered repeatedly beginning 1 hr after SE onset and then four additional times at 24-hr intervals for a total of 5 treatment days, led to a significant reduction in the time until first observed Racine stage 5 seizures (Figure 4B; X^2^=5.691, p=0.0171); i.e., PB-treated rats presented with stage 5 seizures earlier than VEH-treated rats. No dose or time point of administration of EVL administration (Figure 4) led to any significant deviations in the latency to stage 5 seizure versus the respective VEH-treated control cohort (Figure 4D, X^2^=2.280, p=0.131; Figure 4F, X^2^=0.1155, p=0.734). This also includes the head-to-head evaluation of EVL in the original versus expedited protocol (original protocol Figure 4H, X^2^=0.0009090, p>0.97; expedited protocol Figure 4J, X^2^= 0.5971, p>0.44). Thus, EVL treatment did not significantly delay or decrease the time to onset of stage 5 seizures.

## Discussion

In 2016, the NINDS ETSP refocused its *in vivo* approach to evaluate novel treatments for epilepsy to both focus on the identification of agents to treat symptomatic seizures of drugresistant epilepsy and to identify agents that may potentially prevent the development of epileptogenesis or attenuate disease burden in people with epilpesy.^3; 4^ Our present study establishes the proof-of-concept of our screen in a well-established rat model of SE-induced TLE ^21^ using two agents administered by four different treatment designs to formally, rigorously, and blindly evaluate the suitability of this approach. Further, this study reveals our efforts to refine and optimize a testing strategy to provide a moderate-throughput screen for antiepileptogenic agents. While no treatment administered in this study significantly altered the burden of epilepsy, our findings indicate the suitability of this approach to screen for potentially efficacious agents. No known agents can definitively prevent the development of clinical epilepsy and our present negative findings critically inform future antiepileptogenesis trials.

Pharmacological prevention of epilepsy in at-risk individuals has long been considered the “holy grail” of epilepsy therapy development.^1; 2^ There have been clinical studies to prevent the development of epilepsy in post-traumatic brain injury patients using conventional ASDs, including PB, however no such study has yet demonstrated a meaningful effect.^12; 22; 23^ Clinical studies of antiepileptogenesis are incredibly time- and resource-intensive, not to mention sub-optimal because there is no clear timeframe in which epilepsy develops in at-risk individuals. Preclinical models of epileptogenesis are thus essential to identify drug candidates that may prevent epilepsy in at-risk individuals. Neurological insult, including SE, traumatic brain injury, and stroke, are well-known to cause clinical acquired epilepsy, accounting for some 15% of new epilepsy cases.^24; 25^ Our present study evaluated the potential of two mechanistically different agents, PB and EVL, to prevent the development of SRS when administered at three different time points post-SE insult using the well-established rat KA SE model of acquired TLE. Unlike prior studies that have demonstrated a disease-modifying effect of agents in this model,^26; 27^ our study design employed a clinically-relevant interventional approach; namely, sequential evaluation of EVL treatment 1-24 hr after SE onset. Because of the variety of causes of SE, SE is often not able to be treated until well after onset.^28; 29^ When SE is treated in the clinic, interventions are chosen in order to immediately stop the seizure. Our study thus intervened after the neurological SE insult so as to define whether any agent could prevent the development of epilepsy without directly attenuating SE. Whether this design will identify promising antiepileptogenic agents is subject to further scrutiny. In any case, our approach, analytical plan, and interpretation may identify such an agent with the utmost rigor.

EVL and PB are both FDA-approved for the management of epilepsy syndromes; EVL is approved for the treatment of TSC, whereas PB is an ASD approved for monotherapy in diverse seizure types. Limited preclinical studies have assessed the antiepileptogenic potential of both agents. Over 1-month of once-daily PB administration (70 mg/kg, i.p.) beginning 34 minutes after KA-SE insult in PND35 rats did not subsequently prevent the development of SRS and cognitive deficits.^30^ Our present study aligns with these earlier findings using a markedly reduced experimental timeline (5 days of PB treatment). Our study confirms that chronic or subchronic PB is not a valid means to prevent SRS in post-KA SE rats. While Sutula and colleagues demonstrated that PB could prevent the SE-induced damage and modify disease severity when administered during active KA-induced SE, that study administered PB “immediately after” KA administration.^26^ Thus, PB likely attenuated the severity of the SE insult itself, which may have contributed to the observed effects.^26^ Use of a 1 hour delay may suggest that a critical window of PB intervention (i.e. less than 30 min) exists to effectively prevent the development of SRS, likely through the modification of the insult itself.^30^

Similar to PB, EVL did not demonstrate a dose-or time-related antiepileptogenic effect in this model, including a head-to-head evaluation of the original versus expedited protocol designed to improve drug screening throughput. To our knowledge, only one other study has specifically evaluated the preclinical efficacy of EVL using the two-hit mouse model of KA SE-induced TLE.^27^ That study, however, was limited in that EVL (1 mg/kg, i.p.) was administered between the first and second KA bouts, and significantly reduced the latency to Racine stage 5 seizure and attenuated SE severity.^27^ Thus, any observed effects of EVL in that prior study were limited by the initial attenuation of the SE insult. Additionally, there have been conflicting reports regarding the extent to which mTOR inhibition with rapamycin is disease-modifying in various models of acquired epilepsy, ^31-33^ including mouse models of TSC.^34^ Thus, our study suggests through unbiased assessment, that EVL, at the doses and treatment regimen tested, was not antiepileptogenic in the post-KA SE rat model. Our study design provides a robust and unbiased assessment of antiepileptogenic potential *<u>after</u>* SE insult; any effects can be directly attributed to the compound itself, rather than insult modification.

Our present investigations were limited by a number of factors. First, we did not conduct pharmacokinetic assessments of brain concentrations of any compound, nor did we assess the extent to which any of the pharmacological targets (e.g. GABA_A_ receptors or mTOR inhibition) were engaged. In any case, it is entirely possible that our team can perform such bioanalysis. Second, the original design did not include continuous 24/7 vEEG monitoring throughout the 42-day period, but instead broke the observations into two, 2-week sessions separated by a 2-week break. The rationale behind this approach was namely one of logistics. An alternating 2-week on/2-week off/2-week on monitoring protocol can accommodate 24 chronically implanted rats in a single 12-unit recording suite in a standard housing room (10 W x 10 L ft) to conduct long-term studies in a moderate-throughput capacity over roughly 10 weeks. Our original protocol minimized resource burden and increased throughput to conduct long-duration studies. By expediting our study design, we establish a new approach for moderate-throughput drug screening. Future studies will reduce the recording burden based on these presently reported studies to restrict initial screening of novel agents to the monitoring of a single, 2-week recording period from 4-to 6-weeks post SE-insult (expedited). If a future compound is found to modify disease trajectory using this expedited protocol, additional longer duration monitoring can be performed. Finally, our study was limited by the goal to not intervene too early after SE insult so as to clearly dissociate insult-modifying from disease-modifying agents.^35^ By delaying intervention for 1 hour post-SE, we can dissociate any potential disease-modifying effects due to the blockade or reduction in the SE severity itself; in contrast to SE pretreatment studies.^26^ Thus, our design is not without limitations, but we herein offer the epilepsy research community a rigorous, unbiased, and rational screen to identify possible antiepileptogenic agents. Whether our approach will identify any clinical candidates remains to be defined. In any case, this platform supported by the NINDS ETSP offers a critical resource to address the unmet need to prevent epilepsy, as identified by the National Advisory Neurological Disorders and Stroke (NANDS) Council.^3; 4^

## Acknowledgements

We confirm that we have read the Journal’s position on issues involved in ethical publication and affirm that this report is consistent with those guidelines. The authors are grateful for the scientific input of Drs. Brian Klein and John Kehne of the NINDS ETSP. The authors acknowledge the technical support for KA-SE induction of Ms. Stephanie Mizuno. The authors also gratefully recognize the efforts of Dr. Gaëlle Batot for effectively managing the subcontract from the University of Utah. This project has been funded with Federal funds from the National Institute of Neurological Disorders and Stroke, National Institutes of Health, Department of Health and Human Services, under Contract No. HHSN271201600048C (PI KSW) through a subcontract from the University of Utah to the University of Washington.

## Author Contributions

Conception and experimental design: Barker-Haliski, Metcalf, Wilcox, White

Acquisition of experimental data: Knox, Zierath, Koneval, Barker-Haliski

Performed data analysis: all authors

Wrote or contributed to the writing of the manuscript: all authors

