## Supplemental Methods File for "Development of an Antiepileptogenesis Drug Screening Platform: Effects of Everolimus and Phenobarbital"

Supplemental Files.

**Methods.**

*Animals.* All animal experimentation was approved by the University of Washington Institutional Animal Care and Use Committee. Male CD IGS Sprague-Dawley rats (150-200g; Charles-River Laboratories, Wilmington, MA, USA) were housed 5/cage for 1-week to acclimate to the vivarium. After EEG implantation (described below), rats were housed individually in custom Plexiglass cages with corncob bedding, in a temperature-controlled vivarium on a 14:10 light/dark cycle. Animals were permitted *ad libitum* access to irradiated chow (Picolab 5053), filtered water, and enrichment (Nylabones and cardboard tubes). Rats were given a minimum of 24 hours to acclimate to the EEG recording suite prior to all experimentation. Rats were euthanized by CO2 asphyxiation at completion of all in-life studies, in a manner consistent with AVMA guidelines.[1]

To accommodate a target video EEG (vEEG) monitoring group size of n = 12 rats/treatment group (VEH or investigational compound) and potential for SE-induced mortality, a total of 34 animals were surgically implanted 1 week prior to KA SE for each treatment condition (Figure 1). Each study was divided into two cohorts of randomly assigned rats (n = 17/cohort). This allowed for potential KA SE mortality of up to 12-18% (final cohort size of n=14-15 rats); rats that survived the SE insult were then candidates for drug intervention at the relevant time point (Figure 1). Any rat that lost more than 20% of pre-SE body weight within 7 days of SE insult and during the investigational drug or VEH administration period was removed from the study.

*EEG Implant Surgery.* Rats were stereotaxically implanted with a three -prong, cortical recording electrode (MS333/1-A; Plastics One) under ketamine/xylazine anesthesia. All electrodes were implanted posterior to bregma, with the recording electrodes implanted to the right of the sagittal suture line and the ground electrode placed posteriorly left of the suture line. The electrode for each rat was secured to the skull using dental acrylic. Rats were provided post-operative supportive care, including bacon-flavored carprofen (Bio-serv) and allowed to recover for one week in individual housing cages following surgical procedures prior to SE induction.

*Kainic Acid (KA) Model of Status Epilepticus (KA).* Convulsive and electrographic SE was induced by administering KA (i.p.; Tocris catalog 0222 dissolved in sterile saline) via a repeating low-dose protocol, 24 hr after being connected to the EEG monitoring system. All animals received an initial 10 mg/kg dose of KA followed by a 1-hour observation period with subsequent 5 mg/kg doses administered every half-hour until SE onset (defined as two generalized stage 4/5 seizures observed within a 30 min interval and the presence of sustained electrographic high amplitude spiking). The dose of KA was reduced to 2.5 mg/kg if a rat only had a single generalized seizure before the 0.5 hr interval elapsed to minimize potential SE-induced mortality. Behavioral seizure severity and frequency were then scored by trained investigators for 1 hr following SE onset. At the conclusion of a 1-2 hr observation period for all studies, 3 mL of Lactated ringer's (s.c) was provided to each rat to replace any SE-induced fluid loss. Standard rodent chow softened with pediatric electrolyte solution and Napa Nectar hydration gel (Systems Engineering Lab Group, Inc.; Napa, CA USA) were provided for 7 days post-insult as supportive care.

*Investigational Compound and Formulation Vehicles.* The investigational compound, phenobarbital (PB; Sigma Aldrich catalogue #P1636), was formulated for repeated administration in 0.9% saline (Day 1) or 40% hydroxy-propyl beta-cyclodextrin (Day 2-5; Sigma Aldrich catalogue #H107). EVL (MedChem Express # HY-10218) was initially dissolved in 100% EtOH as a 10 mg/mL stock solution and stored at -20C. No solubility or formulation issues were noted with any preparation. For each treatment day, EVL stock was then freshly diluted into a 1 mg/mL dosing solution with a vehicle of 5% PEG-400 (Sigma Aldrich catalogue #06855)/5% Tween-80 (Sigma Aldrich catalogue #P-1754) in 0.9% saline. Because there were different formulation vehicles for PB and EVL, it was necessary to have an independent VEH-treated KA-SE control group for each investigational compound.

Each investigational compound was administered for five (PB and EVL 3 mg/kg) or seven (EVL 2 mg/kg) consecutive days after SE onset (Figure 1). For each study, animals were randomized to their treatment group based on time of SE onset. Investigators were blinded to treatment group (n=12/ treatment group). PB (60 mg/kg) or its VEH was administered by the intraperitoneal (i.p.) route on Day 1 to minimize any potential for trauma during the sustained KA-induced SE, and then by the subcutaneous (s.c.) route on Days 2-5 post-SE. PB or VEH administration was started 1 hr after onset of behavioral SE, with SE confirmed by sustained electrographic spiking. The dose and time of administration of PB was based on prior studies demonstrating no disease-modifying effect of PB when administered >30 min after SE onset.[2]

Supplemental Files.

EVL (2 mg/kg) or its VEH was administered by the oral (p.o.) route for 7 days beginning 24 hr after onset of behavioral and electrographic SE (Figure 1). EVL (3 mg/kg) or its VEH was administered p.o. route for 5 days beginning 2 hrs after SE onset (Figure 1). The doses of EVL and time points of administration were selected based on prior studies to suggest that rapamycin, an agent with limited brain bioavailability, could modify disease severity in a post-TBI model of epileptogenesis in mice.[3] No other pharmacological intervention was administered during the post-KA SE period or up to 42 days post-insult.

*SE and Spontaneous Seizure Scoring.* Both the initial SE and resultant SRS were scored according to the Racine scale, with generalized seizures scored highest (stages 4 and 5). Additionally, during the chronic monitoring period, severe and sustained “popcorn” seizures that occurred after stage 5 bilateral forelimb clonus, rearing, and falling were scored as stage 6. Rats were monitored via video EEG (vEEG) for up to 6 weeks post-SE onset. In the original study protocol, synced video and EEG data were recorded continuously in two, 14-day bins (weeks 0-2, weeks 4-6), separated by two weeks of non-recording (weeks 2-4). In the “expedited” protocol, synced video and EEG data were recorded continuously in a single 14-day session during 4-6 weeks post-SE).

*vEEG monitoring of spontaneous seizures*. Video-EEG was recorded using a customized data acquisition system with an MP160 and EEC100 specify tethered / wireless system (Biopac, Goleta, CA) and 3-channel electrodes (InVivo1).[4] Following each daily recording, the vEEG files for all rats were manually reviewed in custom vEEG playback software by an experienced investigator blinded to treatment condition. The files were played back at 2x-10x speed and visually scanned for patterns of high frequency electrographic spiking followed by a depression in electrographic activity that are typical of SRS. When a rat’s EEG channel displayed a possible electrographic seizure-like event, it was selected, and the paired video was reviewed in real time to confirm whether a convulsive seizure occurred. The convulsive seizure was scored on the Racine seizure scale.[5] Briefly, behavioral seizures were scored as: Stage 1 - mouth and facial clonus; Stage 2 - Stage 1 activity + head bobbing/nodding; Stage 3 - Stage 2 activity + unilateral forelimb clonus; Stage 4 - Stage 3 activity + bilateral forelimb clonus and rearing up onto hindlimbs; Stage 5 = Stage 4 activity + falling over. If a convulsive seizure did not occur, the event was classified as either a false-positive (e.g. head scratching, chewing, grooming) or defined by any notable changes in behavior, such as prolonged freezing while awake, that was marked as a non-convulsive event. Upon completion of each continuous 2-week recording session, the seizure scores of all rats in each study cohort that occurred during the entire recording session were reviewed by a secondary investigator to confirm seizures and severity. The cumulative seizure burden for each animal was then used to assess the efficacy of each investigational compound (PB 60 mg/kg, EVL 2 mg/kg, or EVL 3 mg/kg) versus its respective vehicle.

The seizure burden of each rat within each 2-week recording session was quantified, which is a measure of the Racine stage x frequency of each Racine stage event. Cumulative seizure burden was also determined, which is the summation of all observed events during each of the two recording sessions. Cumulative seizure burden can be a useful index of disease progression during the entire study period.[6; 7] The cumulative seizure burden was quantified up to 42 days post-SE to define the extent to which an intervention altered the trajectory of disease progression.

*Statistics.* The change in body weight for each treatment group within each study was analyzed by repeat measures ANOVA (time post-SE x treatment). The average number of seizures, average seizure burden, average seizure severity, and average seizure frequency was binned for each 2-week recording session and analyzed by repeat measures ANOVA (time post-SE x treatment) across the two recording sessions. The latency to stage 5 seizure was analyzed by Kaplan-Meier survival plot. The cumulative seizure burden throughout the study period (not per binned sessions), which is a continuous sum of each day's seizure scores added across days in each recording session, was analyzed by repeat measures ANOVA. For all statistical analyses, data was analyzed with GraphPad v7.0 or later with $p<0.05$ considered statistically significant.

**Barker-Haliski, M. et al. -** ***Development of an Antiepileptogenesis Drug Screening Platform: Effects of Everolimus and Phenobarbital***

Supplemental Files.

Supplemental Figures.

Supplemental Figure 1. Effects of pharmacological intervention on change in body weight from pre-kainic acid (KA) status epilepticus (SE)-induced body weight (baseline) in the original disease-modification protocol. The original protocol design included EEG electrode implants 7-10 days prior to systemic low-dose KA administration to adult male Sprague Dawley rats to induce SE. Following the confirmed onset of sustained KA-induced SE (with electrographic and behavioral seizures), rats were administered either A) phenobarbital (PB; 60 mg/kg), B) everolimus (EVL; 2 mg/kg, p.o.), or C) EVL (3 mg/kg, p.o.) beginning no earlier than 1 hour post-SE onset. For all treatment conditions, there was a significant main effect of time post-SE induction body weight change from Day 0 (SE induction day). All rats in all treatment groups also demonstrated significant body weight loss due to the KA-induced SE insult (change from Day 0 to Day 1 post-SE). There were no significant differences in body weight change between VEH- and investigational drug-treated rats for any treatment condition (PB or EVL at either dose).

Supplemental Files.

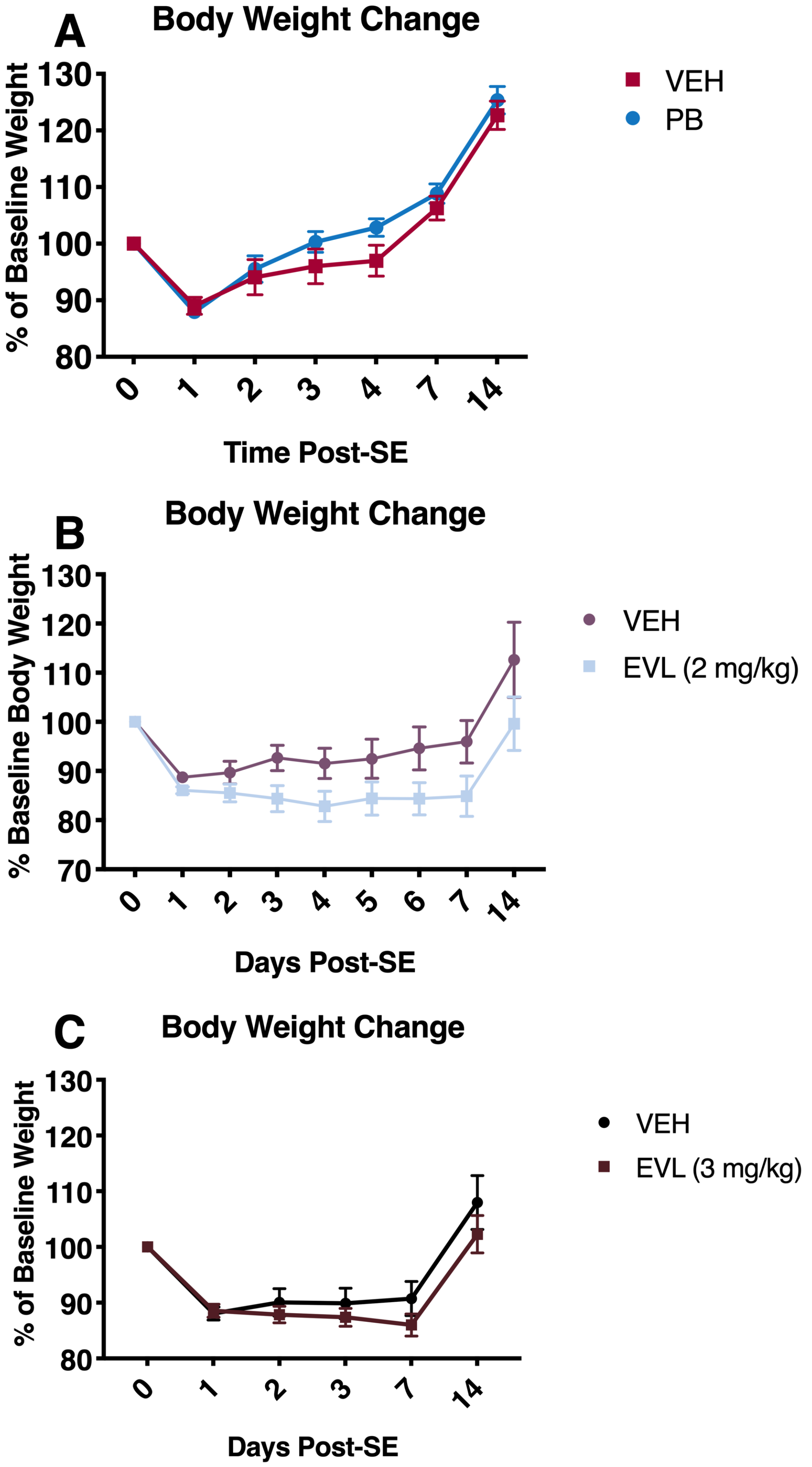

**Barker-Haliski, M. et al. - *Development of an Antiepileptogenesis Drug Screening Platform: Effects of Everolimus and Phenobarbital***

Supplemental Files.

Supplemental Figure 2. Effects of pharmacological intervention on change in body weight from pre-kainic acid body weight (baseline) in the original versus expedited disease-modification protocols. The original protocol included EEG electrode implants 7-10 days prior to systemic low-dose KA administration to male Sprague Dawley rats, whereas the expedited protocol conducted EEG implants 7 days after KA-induced SE to minimize the numbers of animals undergoing surgery. Further, the original protocol included vEEG monitoring from 0-2 and 4-6 weeks post-SE, whereas the expedited protocol only included vEEG monitoring from 4-6 weeks post-SE. In both protocol designs, rats received EVL (3 mg/kg, PO, q.d.) beginning 1-hour post-onset of KA-induced SE. For both protocol designs, there was a significant main effect of time post-SE induction body weight change from Day 0 (SE induction day; A) $F(1.768, 45.98) = 37.66$, $p<0.0001$; B) $F(1.806, 48.75) = 65.61$, $p<0.0001$). All rats in all treatment groups also demonstrated significant body weight loss due to the KA-induced SE insult (change from Day 0 to Day 1 post-SE). However, there were no significant differences in body weight change between VEH- and EVL-treated rats in either protocol design (original or expedited). Thus, the expedited protocol design (B) is not associated with any marked differences in capacity to detect changes in SE-induced body weight change versus our original disease-modification protocol (A).

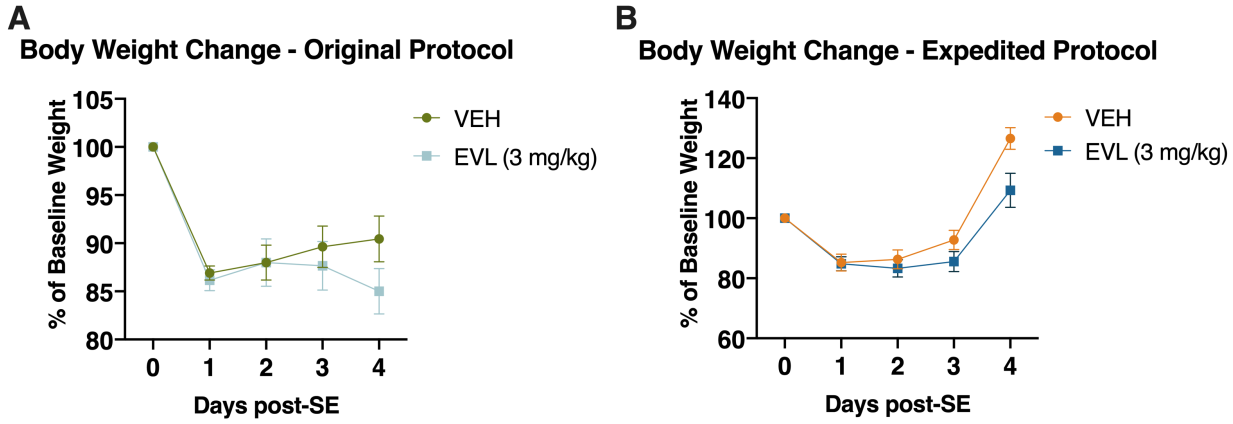

**Barker-Haliski, M. et al. -** ***Development of an Antiepileptogenesis Drug Screening Platform: Effects of Everolimus and Phenobarbital***

Supplemental Files.
